# Theory of neural coding predicts an upper bound on estimates of memory variability

**DOI:** 10.1101/793430

**Authors:** Robert Taylor, Paul M Bays

## Abstract

Observers reproducing elementary visual features from memory after a short delay produce errors consistent with the encoding-decoding properties of neural populations. While inspired by electrophysiological observations of sensory neurons in cortex, the population coding account of these errors is based on a mathematical idealization of neural response functions that abstracts away most of the heterogeneity and complexity of real neuronal populations. Here we examine a more physiologically grounded model based on the tuning of a large set of neurons recorded in macaque V1, and show that key predictions of the idealized model are preserved. Both models predict long-tailed distributions of error when memory resources are taxed, as observed empirically in behavioral experiments and commonly approximated with a mixture of normal and uniform error components. Specifically, for an idealized homogeneous neural population, the width of the fitted normal distribution cannot exceed the average tuning width of the component neurons, and this also holds to a good approximation for more biologically realistic populations. Examining eight published studies of orientation recall, we find a consistent pattern of results suggestive of a median tuning width of approximately 20 degrees, which compares well with neurophysiological observations. The finding that estimates of variability obtained by the normal-plus-uniform mixture method are bounded from above leads us to reevaluate previous studies that interpreted a saturation in width of the normal component as evidence for fundamental limits on the precision of perception, working memory and long-term memory.

## Introduction

The continuous report task (Wilken & Ma, 2004; Prinzmetal, Amiri, Allen, & Edwards, 1998) provides a means of measuring the fidelity with which visual information can be retained in memory. The general procedure first presents observers with a set of stimuli differing in an elementary visual feature, such as color hue or orientation, which they are required to remember during a short retention interval. At test a single target item is identified (e.g., with a cue at its previous location) and observers must reproduce the corresponding memorized feature value via an analogue response method (e.g., clicking on a color wheel). Internal noise ensures that there is some degree of error in observers’ estimates of the target feature, and across trials the dispersion of these errors provides a metric of memory precision. Numerous studies testing memory for a wide variety of stimulus features have shown that the variability in recall errors (as measured by, e.g., their standard deviation [s.d.]) increases smoothly and continuously with set size (the number of items in the memory array; Bays, Catalao, & Husain, 2009; van den Berg, Shin, Chou, George, & Ma, 2012; van den Berg, Awh, & Ma, 2014).

While the s.d. of errors provides a concise summary of recall fidelity in a particular experimental condition, attempts have been made to further distinguish different types of error that might contribute to the pattern of responses. One popular method, the normal-plus-uniform model (W. Zhang & Luck, 2008), statistically decomposes responses into two components, with the intention of distinguishing responses based on memory for the target item from random guesses. This method assumes that responses resulting from memory of the target item will have a von Mises distribution (a circular analogue to the normal) centered on the true feature value, whereas other responses will be uniformly distributed in feature space. According to this model, it is the standard deviation of just the fitted von Mises component of the error distribution that measures the precision with which items are stored, whereas the mixing proportion of the fitted von Mises and uniform components reflects the probability of a tested item being in memory at all.

A subsequent study (Bays et al., 2009) found that the uniform component of the normal-plus-uniform fit captured many responses that were in fact distributed around the features of other, non-target items in the memory array, and proposed adding a third component to the model to capture these “swap errors”. The relative proportion of swap errors may be estimated using either parametric methods (Bays et al., 2009; van den Berg et al., 2014; see also Rerko, Oberauer, & Lin, 2014), or non-parametric approaches (Bays, 2016a). These errors have proved an important source of information for understanding how multiple features of an object are linked or “bound” together in memory (Schneegans & Bays, 2018). However, in this study we focus primarily on interpreting the von Mises component of the normal-plus-uniform model, so for the present purposes it is sufficient to say the other component is uniformly distributed *with respect to the target item*.

Bays (2014) noted that working memory error distributions corresponded very closely with those predicted by a simple encoding-decoding model of working memory. This model is based on the principles of population coding (Pouget, Dayan, & Zemel, 2003, 2000), and assumes that visual feature information is first encoded in the activation of feature-selective neurons, and subsequently reconstructed from the persisting (or reinstated) spiking activity of the same cells. This study further found that the effects of set size on recall error could be parsimoniously explained if the total activity of the neural population served as a limited resource, shared out between memory items (i.e. if the activity was normalized, see Bays, 2015, for neural evidence and possible mechanisms). As a consequence, for larger memory arrays each item is represented with fewer spikes, so the model predicts that item precision declines as set size increases. Formal model comparison has shown that the neural model provides a better account of recall errors than models based on a mixture of remembered and guessing states (Bays, 2014; Taylor & Bays, 2018; van den Berg, Yoo, & Ma, 2017). The neural model also quantitatively reproduces the effects of predictive (Bays, 2014) and retrospective cues on recall (Bays & Taylor, 2018) and accurately predicts both the frequency of swap errors and which non-targets are likely to be reported in place of the target (Schneegans & Bays, 2017).

While the population coding account has been very successful at reproducing patterns of error, it is based on a mathematical idealization of neural response functions that abstracts away most of the heterogeneity and complexity of real neuronal populations. We therefore set out to examine whether a neural population that better reflects the tuning characteristics of visual cortical neurons can reproduce benchmark behavioral results. To anticipate our results, we found that key predictions of the idealized model are indeed preserved. In particular, physiologically derived populations also predict long-tailed distributions of error at lower levels of population activity, as observed empirically at higher set sizes and for items with lower priority in memory.

We subsequently investigated the consequences of fitting the normal-plus-uniform mixture model to recall distributions generated from each neural population. Irrespective of their theoretical basis, the mixture parameters can be viewed as descriptive statistics, concisely summarizing the patterns of response errors predicted by the population models. Our central finding was that the estimated width of the normal mixture component (which measures the “central peak” in error distributions) cannot exceed the average width of the tuning functions in the underlying neural population. We show that results of previous recall experiments support the existence of such an upper bound, and that the bound is broadly consistent with typical tuning widths recorded in electrophysiological studies.

These results have several important implications. First, they provide validation for the population coding model by confirming a correspondence between behavioral recall performance and observable properties of the underlying neural system – while the comparison is indirect at this stage, technical advances should permit increasingly robust tests of this correspondence in future. Second, they place an alternative, theory-based interpretation on the parameters obtained from the normal-plus-uniform fit. Hundreds of experimental studies have reported and interpreted their results on the assumption that this fit accurately distinguishes random guesses from memory-based responses. We show that these previous results can be re-interpreted, rather than simply discounted, in light of the population coding account, providing valuable information about the working memory system and its relation to perception and long-term memory. Finally, our numerical simulations and the analytical results provided in the Appendix provide some insight into how mechanisms built on continuous-valued functions can produce behavior superficially suggestive of dichotomous (all-or-nothing) states.

## Methods

### Population Coding Model

Population coding provides a mechanistic description of how visual inputs are registered by feature-selective neurons (Ma, Beck, Latham, & Pouget, 2006; Pouget et al., 2003, 2000). The principle is that a visual feature value – e.g., the orientation of a contrast edge – evokes a pattern of noisy activity in a population of neurons that imperfectly encodes that feature value. As a model of memory, the assumption is that this activity can be maintained, or restored after a delay, at which point the original feature value can be decoded from the joint population activity. The inherent variability in neural spiking (often equated with a Poisson process) ensures that decoded values will be probabilistically distributed around the original stimulus value. The width and shape of this error distribution is principally determined by two parameters of the neural population: the summed activation (or gain) of the population and the tuning specificity of the cells.

We conducted simulation studies to understand the consequences of fitting the normal-plus-uniform model to the distributions of error predicted by population coding. We considered two different types of neural population: our first simulation was based on an idealized population which has been used previously to fit empirical response distributions and for which error distributions can be obtained quite directly through mathematical analysis (for details, see Bays, 2016b). In our second study, we considered a more biologically realistic population with tuning properties drawn from real neurons, where due to the model’s complexity error distributions can only be obtained through numerical simulation. We describe the details for each next.

#### Idealized neural population

In this population, neurons had identical, von Mises tuning functions, evenly spaced throughout the input feature dimension, with no baseline activity (see top left panel in Fig. 2). Additionally, each neuron’s spiking activity was independent of all the other neurons’ activities. Formally, for *M* neurons, the firing rate of the *i^th^* neuron with preferred value *φ_i_* in response to a stimulus value *θ* was given by:

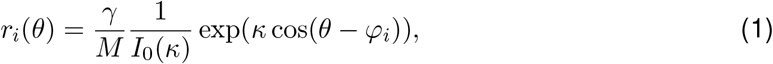

where *γ* is the population gain, *κ* is the tuning specificity, and *I*_0_(·) is the modified Bessel function of the first kind with order zero. Spikes were generated by a Poisson process, and decoded feature values were obtained via maximum likelihood (for full details, see Bays, 2014).

The gain parameter, *γ*, controls how responsive the population is to visual stimulation and determines how many spikes are available for decoding feature values. As the population gain increases, the model can encode visual information more reliably which typically produces smaller errors between the decoded and veridical feature values. The tuning specificity parameter, *κ*, defines the set of input values that elicit a response from each neuron in the population. Broadly tuned neurons (small *κ*) respond to a wider range of input values whereas narrower functions (large *κ*) have a more selective response. Bays (2016b) previously showed that, for large *M*, error distributions can be approximated by a von Mises random walk, where the number of steps is determined by the total spike count observed during the decoding window. Conveniently, this permits the error probability density function, *f_neural_*(*ε*; *γ*, *κ*), to be straightforwardly obtained for any values of the model parameters (see Bays, 2016b, for derivations; code available at www.bayslab.com/code/JN14/).

We generated predicted error density functions by parametrically altering the tuning and gain parameters. A parameter grid was instantiated that contained four *κ* values, {2, 4, 8, 16}, and 40 *γ* values, logarithmically spaced between 10^−1.5^ and 10^2^. For each parameter combination we estimated normal-plus-uniform model parameters by minimizing the sum of squared errors (SSE) between the neural error distribution and the mixture model error distribution (see Eq. 6), evaluated at 10^3^ evenly-spaced points on the circle:

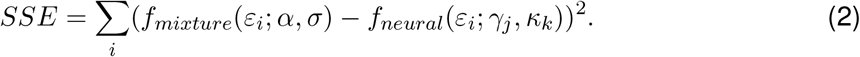

#### Biological neural population

Here we describe simulations of a neural population based on real tuning characteristics observed in visual cortex. We focused on the properties of orientation-selective neurons in area V1 (primary visual cortex) as a canonical example of population coding. To our knowledge no study in humans has mapped out individual orientation tuning functions of V1 neurons. However, a study in non-human primates conducted by Ecker et al. (2010) provides suitable data, based on electrophysiological recordings of 408 neurons in primary visual cortices of two macaques viewing oriented sine wave gratings. Tuning characteristics of each recorded neuron, including baseline activity, amplitude and tuning specificity, were publicly released alongside code from (Ecker, Berens, Tolias, & Bethge, 2011; downloaded from http://bethgelab.org/code/ecker2011/). This data provided the basis for our simulations of heterogeneous population coding.

In the heterogeneous neural population every component neuron possessed an individually parameterized tuning function (see top right panel in Fig 2 for examples). Given a specific input value, *θ*, the mean response of the *i^th^* of *M* neurons was described using the following general expression for a von Mises tuning function with baseline activity,

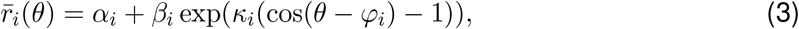

where *α_i_* is the neuron’s baseline activation level, *β_i_* its amplitude, and *κ_i_* its tuning specificity. We generated heterogeneous populations by drawing tuning parameters randomly with replacement from the pool of neurons characterized by Ecker et al. (2010). As the summary dataset made available by these authors did not include preferred orientation values, *φ_i_*, these were randomly drawn from a uniform distribution on the circle. The randomly drawn tuning functions were then scaled to produce a median tuning specificity of 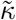 and an expected population gain of 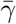.

We further introduced short-range pairwise correlations between neurons. We modeled correlation in the activity between the *i^th^* and *j^th^* neuron as an increasing function of the similarity between the neurons’ preferred input values (as in Sompolinsky, Yoon, Kang, & Shamir, 2001):

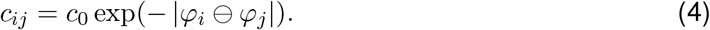

Small positive neural correlations are found throughout cortex though estimates of their magnitude vary considerably: for instance, estimates of correlations between neurons in V1 range from 0.01 to 0.26 (Cohen & Kohn, 2011). For our simulations we fixed *c*_0_ at 0.2, choosing a value at the upper end of existing estimates in order to more clearly observe any consequences of correlations for the decoded error distributions.

We instantiated a parameter grid that contained four values of the median tuning specificity, 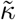, {2, 4, 8, 16}, and 20 population gain 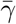 values logarithmically spaced between 10^−2^ and 10^3^.

The predicted error density functions cannot be obtained via the method described by Bays (2016b) and so must instead be approximated via Monte Carlo simulation. Furthermore, simulating correlated Poisson processes is computationally very demanding (e.g. Macke, Berens, Ecker, Tolias, & Bethge, 2009), so we used a Gaussian approximation to Poisson, as in e.g. Schneegans and Bays (2017). Specifically, samples of population activity, ***r***(*θ*), were generated as draws from a multivariate normal distribution 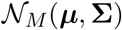, with mean equal to the neurons’ mean firing rates, 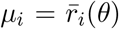, and covariance, 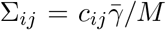, with the result that the variance in total population activity scaled with the mean as for Poisson noise.

For each parameter combination 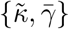, we generated 10^6^ samples of activity from each of 192 randomly drawn heterogeneous populations each comprising 1000 neurons, using the Cholesky decomposition method for generating correlated random variables. Each sample was based on a different stimulus value chosen randomly from a uniform distribution on the circle. Estimation of the encoded stimulus was based on maximum likelihood (ML) decoding of the joint population activity, where we assumed that the decoder was not aware of the correlations in the population. Specifically, we numerically maximized the log likelihood for uncorrelated activity,

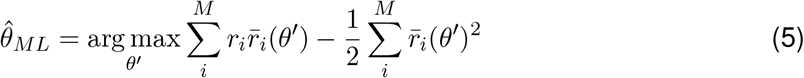

based on discretizing the circular stimulus space into 100 equally spaced bins. We then computed a histogram probability density estimate of the error in the ML estimate, based on the same bins, and collapsing over all samples and simulated populations. Finally, we fit the normal-plus-uniform model to the histogram estimate, as described above for the idealized population.

### Normal-plus-Uniform Model

The normal-plus-uniform model of recall errors (W. Zhang & Luck, 2008) specifies a probabilistic mixture of two distributions: one von Mises centered on the true value of the stimulus to be recalled, and one uniformly distributed across all possible feature values. The probability density function is,

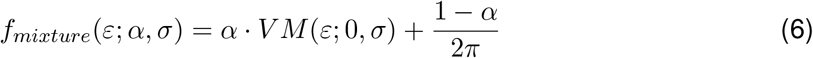

where 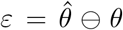 denotes the angular deviation between the target, *θ*, and reported feature value, 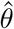, on the circle; *VM*(*ε*; 0, *σ*) denotes the probability density function of a zero-centered von Mises distribution with circular standard deviation *σ*, evaluated at *ε*; and *α* is the mixing parameter determining the proportion of errors drawn from the von Mises distribution. Both *α* and *σ* are free parameters that must be estimated from data.

### Experimental Data

We fit both the idealized population coding model and the normal-plus-uniform model to seven existing orientation reproduction datasets, all either already in the public domain or sourced from our own lab. Some methodological details varied between studies (e.g., whether masking was used, presentation time, retention interval, etc), but the general experimental protocol remained the same: participants studied arrays containing a variable number of oriented stimuli and were subsequently required to reproduce the orientation of a randomly selected target using an analogue response device. Study details are laid out in Table 1 (note that Study 1 manipulated presentation duration in order to assess the temporal dynamics of feature encoding; the model fits presented here were obtained by collapsing across durations. For all further details the reader is referred to the Methods sections of the original studies).

**Table 1.**
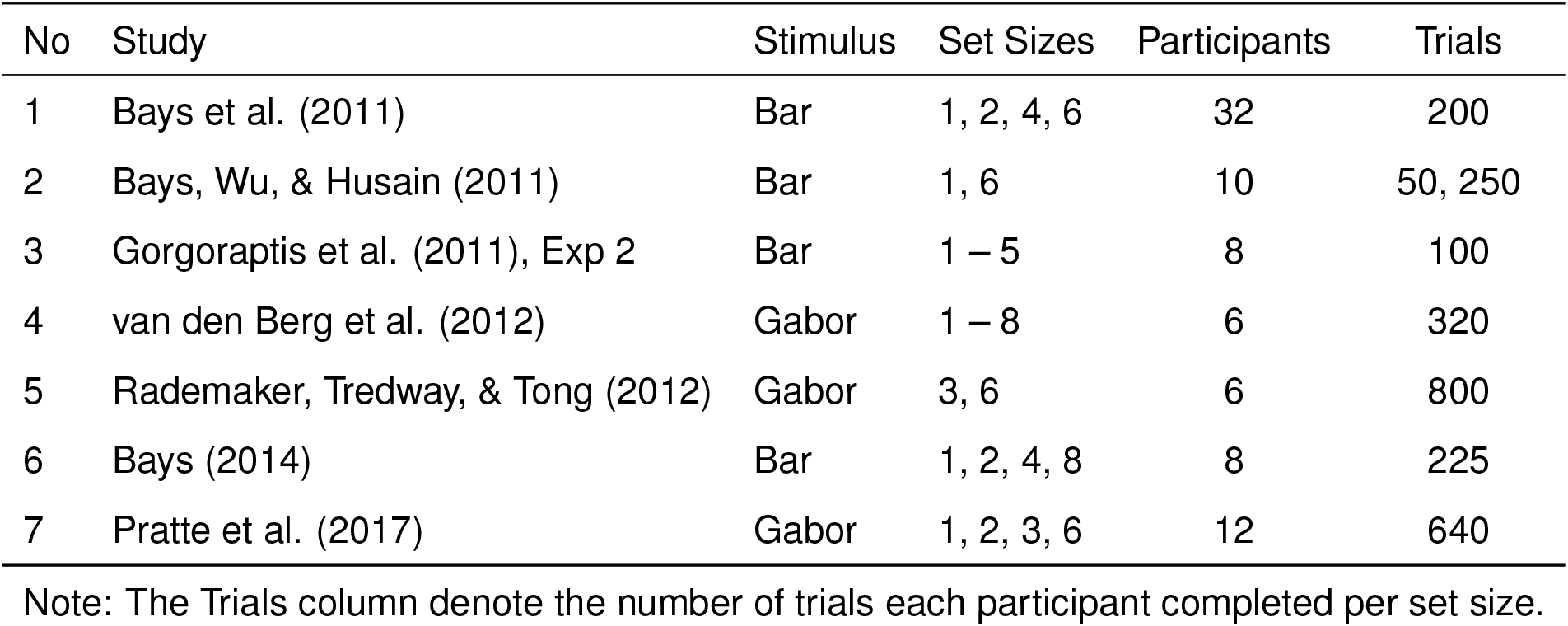
Methodological details of the orientation recall datasets supporting the normal-plus-uniform model fits displayed in Figure 2.

| No | Study | Stimulus | Set Sizes | Participants | Trials |
| --- | --- | --- | --- | --- | --- |
| 1 | Bays et al. (2011) | Bar | 1, 2, 4, 6 | 32 | 200 |
| 2 | Bays, Wu, & Husain (2011) | Bar | 1, 6 | 10 | 50, 250 |
| 3 | Gorgoraptis et al. (2011), Exp 2 | Bar | 1 – 5 | 8 | 100 |
| 4 | van den Berg et al. (2012) | Gabor | 1 – 8 | 6 | 320 |
| 5 | Rademaker, Tredway, & Tong (2012) | Gabor | 3, 6 | 6 | 800 |
| 6 | Bays (2014) | Bar | 1, 2, 4, 8 | 8 | 225 |
| 7 | Pratte et al. (2017) | Gabor | 1, 2, 3, 6 | 12 | 640 |

Note: The Trials column denote the number of trials each participant completed per set size.

For the idealized population coding model, parameters were estimated separately for each participant, with the assumption that gain decreases in inverse proportion to set size (*γ* ∝ 1/*N*, i.e. equal allocation of neural resource as in Bays, 2014; code for fitting the idealized population coding model is available at www.bayslab.com/code/JN14/). For the normal-plus-uniform model, parameters were estimated separately for each participant and set size (code available at www.bayslab.com/code/JV10/). To generate the curve in Fig. 4 we fit the participant-averaged von Mises s.d. parameters from the normal-plus-uniform fit, pooled across experiments, with an exponential saturation function,

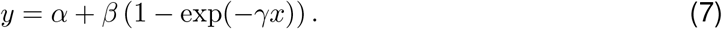

The asymptote of the curve was calculated as *α* + *β*.

Note that there is possibility for confusion when dealing with orientation data, because the range of possible orientations spans 180°, yet this space has the topology of a circle, and circular statistics are naturally conducted on an interval spanning 360°. In this paper, we report orientation values in degrees of the original orientation space, which for clarity we write e.g. 10° [180°]. In contrast, when reporting non-orientation results we do so in degrees on the circle, writing e.g. 10° [360°].

We further examined parameters of the normal-plus-uniform model obtained in four previous studies, all of which tested memory for color. Data from W. Zhang and Luck (2008) is already in the public domain, so we estimated model parameters in the same way as for the orientation experiments. For the remaining studies, individual trial data was not available, so we reproduce here the model parameters reported in the text and/or figures of each paper (Brady, Konkle, Gill, Oliva, & Alvarez, 2013; Asplund, Fougnie, Zughni, Martin, & Marois, 2014; W. Zhang & Luck, 2009).

## Results

We first simulated the reconstruction of an abstract circular stimulus value from activity of one of two classes of neural population. In an idealized population (illustrated in Fig 1a), every neuron’s tuning is described by an identical von Mises function, with neurons differing only in their preferred feature values (the peaks of the tuning functions), which span the feature space with uniform density. Fig 1c shows how the distribution of error in decoding a feature value stored in this idealized neural population (Fig 1a) changes with the mean activity level in the population (gain, *γ*). As reported in previous work (e.g. Bays, 2014), the error distributions deviate from normality, this discrepancy becoming particularly salient at lower activity levels (e.g. blue curve in Fig 1a) where the error distributions grow increasingly long-tailed.

**Figure 1.**
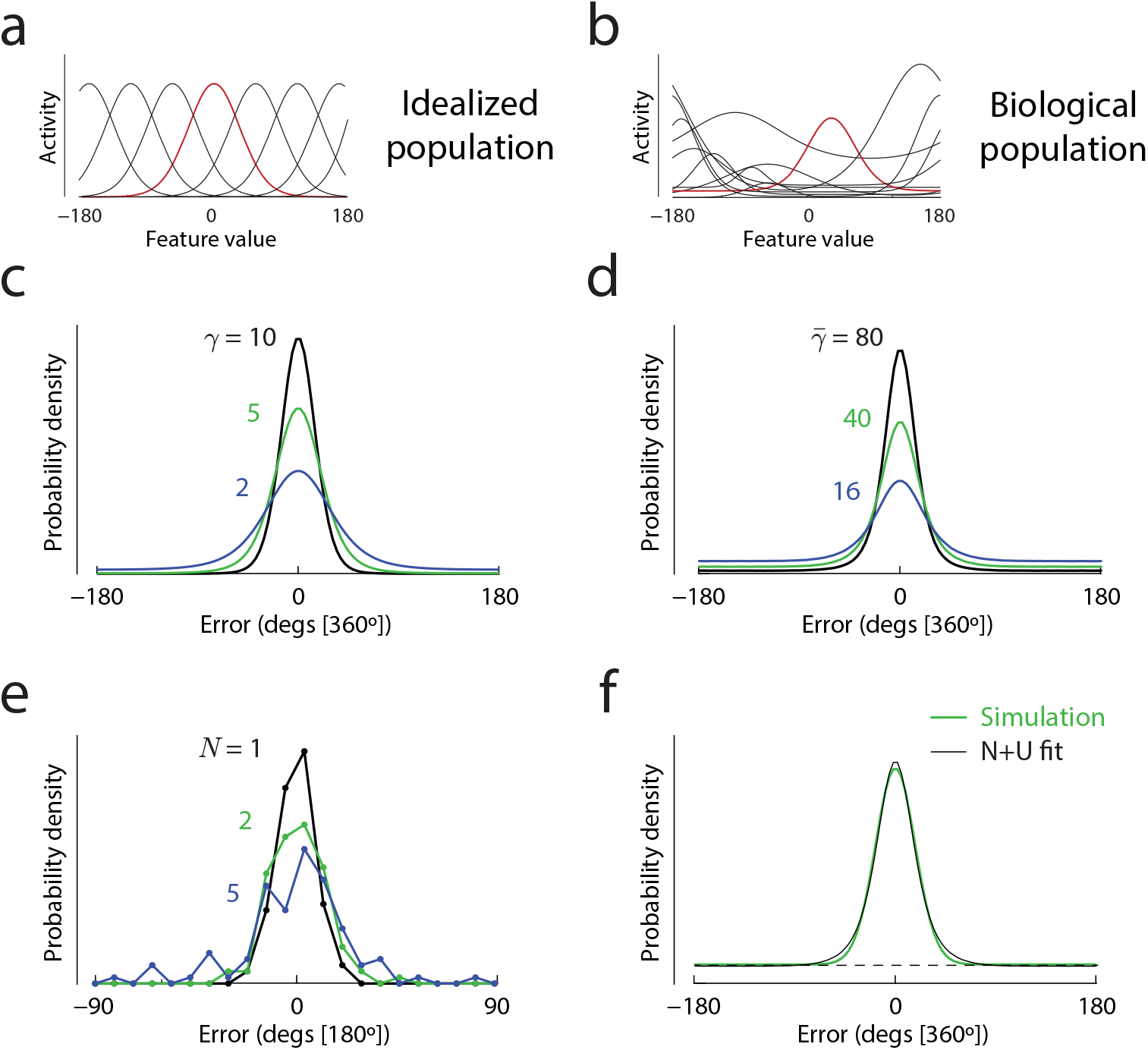
Overview of models and methods. (a & b) Examples of tuning functions in an idealized homogeneous neural population (a) and in a heterogeneous population based on electrophysiological recordings from macaque V1 (b). For each population, the tuning function of one example neuron is highlighted. (c & d) Predicted distributions of error in decoding an abstract feature value from activity of the idealized (c) or biological (d) population. Different colors correspond to different amplitudes of population activity, with higher mean activity levels in the biological population chosen to roughly equate error variability with the idealized model. For this illustration the idealized population had tuning width (*κ* = 2, ~ 49° [360°]). The median tuning width in the biological population was 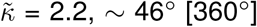. (e) Example distributions of errors made by a single illustrative participant in an orientation reproduction task (Study 3 in Table 1) at three different set sizes (*N*, colors). (f) Approximation of a model-predicted error distribution (green; biological population with 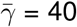) with a normal-plus-uniform mixture (black).

Fig 1b illustrates tuning of a sample of real orientation-selective neurons recorded in macaque V1 by Ecker et al. (2010). While the tuning of each neuron was again approximated by a von Mises function, neurons differed greatly in their tuning width and amplitude, as well as their level of baseline activity. For the purposes of simulation, we generated “biological” populations by randomly sampling tuning parameters from this data set. Based on results from other electrophysiological studies, we further introduced correlated noise into the neurons’ activity (see Methods for full details). Fig 1d shows the error distributions generated by decoding of feature values from a biological population. With appropriate choice of mean activity (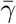; higher than in the idealized model) we observe considerable qualitative similarity between these predictions and those of the idealized model. In particular the presence of long tails at lower activity levels is preserved.

Fig 1e plots distributions of errors made by a representative participant in a typical continuous report experiment, testing recall of one orientation from a display of *N* oriented stimuli. At larger set sizes, the long tails evident in both simulated datasets are visible here too. This aspect of recall errors has often been interpreted with respect to fitted parameters of a normal-plus-uniform mixture model. While the psychological interpretation of the mixture model components is debated, the parameters (the von Mises s.d. and mixing proportion) can be viewed as descriptive or summary statistics capturing key aspects of the error distributions. Fig 1f shows how a mixture of normal and uniform error components (black line) approximates an error distribution obtained by simulation of a biological neural population. In the next section we take advantage of this method to quantitatively compare the predictions of the two neural models.

### Comparing simulations of idealized and biological populations

Fig 2 shows results of fitting the normal-plus-uniform model to the error distributions derived from each neural population. Considering first the standard deviation of the fitted von Mises (i.e. normal) component, for an idealized population of neurons (Fig 2c) we observed that, as the total activity in the population decreased (moving from left to right on the x-axis) the von Mises s.d. initially increased rapidly but then saturated, approaching an asymptotic upper bound. The asymptotic value coincided exactly with the standard deviation of the neural tuning functions, indicated by the dotted lines (different colored lines correspond to populations with different tuning widths).

**Figure 2.**
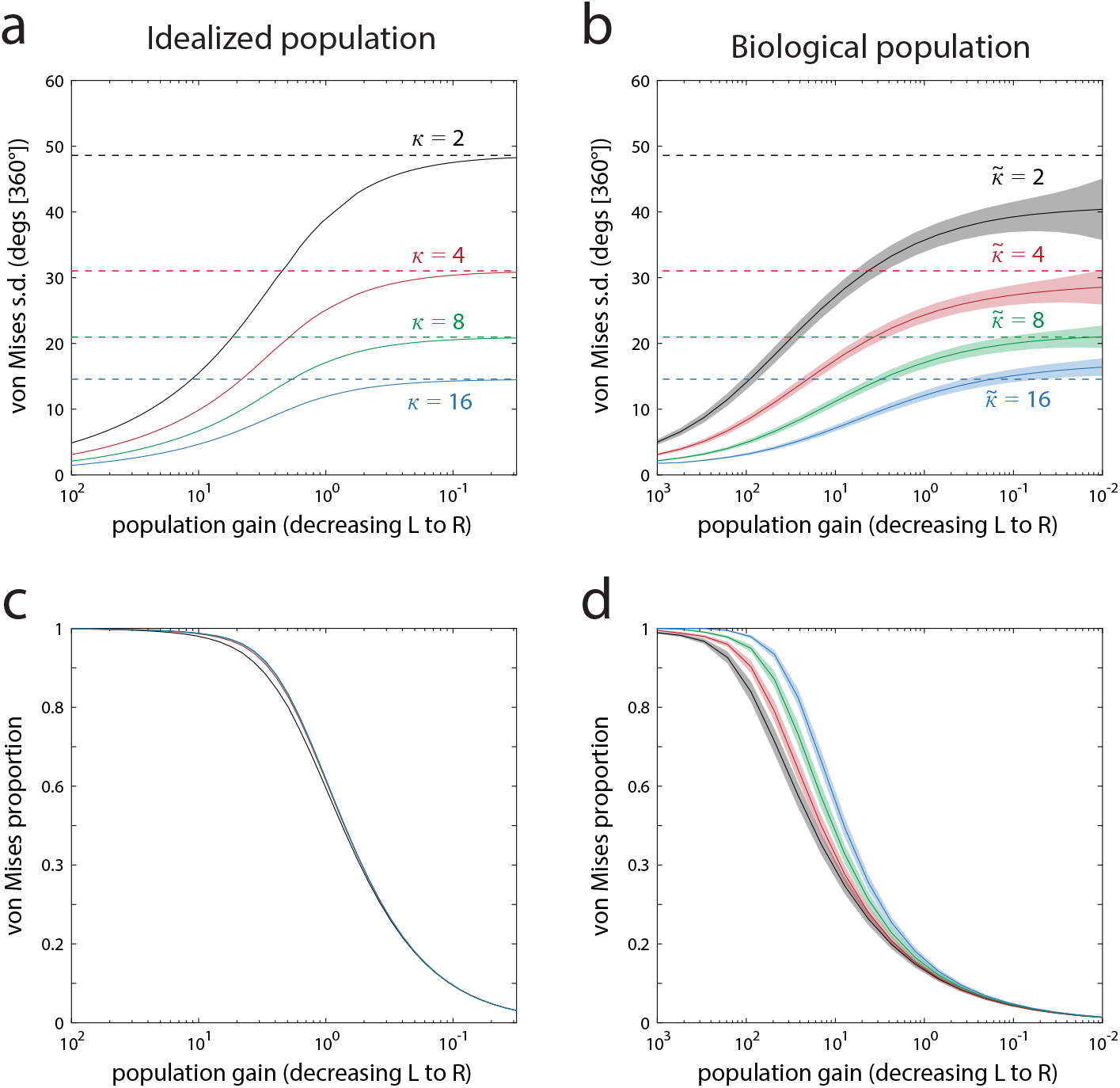
Results of fitting the normal-plus-uniform model to errors resulting from population coding. (a & b) Parameter values for the s.d. of the von Mises component obtained by fitting the normal-plus-uniform model to errors generated from the idealized population (a) and biological population (b). Population gain is plotted on a log axis, decreasing from left to right. Solid lines show fitted parameter values for different neural tuning widths (colors; larger *κ* values correspond to narrower tuning functions; for the biological population 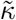 indicates the median tuning width). Dashed lines indicate the s.d. corresponding to each tuning width. Shaded areas in (d) indicate ±1 s.d. of parameter estimates across simulation repetitions. Note that for both idealized and biological populations, as gain decreases the von Mises s.d. parameter approaches an upper bound corresponding (approximately in the biological case) to the average tuning width in the population. (c & d) Parameter values for the mixing proportion of the von Mises component of the normal-plus-uniform fit. Note that, while the s.d. parameter saturates at its upper bound, the mixing proportion continues to fall towards zero as gain decreases.

For a homogeneous population of neurons without baseline activity, this asymptotic behavior is expected, and we have included a mathematical explanation of this result in the Appendix. A simple intuition is that a single spike generated by a tuned neuron narrows down the possible values of the input stimulus to a range equal to the width of its tuning function. In the limit, as the gain of the population approaches zero, each decoded value will be based on either one spike or none. With one spike, the optimal estimate is simply the preferred value of the neuron that generated the spike; for zero spikes, all stimulus values are equally probable. So, assuming approximately von Mises tuning, the distribution of error approaches a mixture of a von Mises centered on the stimulus value with width equal to the underlying tuning width, corresponding to the one-spike state, and a uniform distribution corresponding to the zero-spike state.

Fig 2d shows corresponding results for the population based on V1 electrophysiology. Despite the heterogeneity in tuning and correlations in activity, the pattern of changes to model parameters is remarkably similar to the idealized population. The von Mises s.d. is again found to saturate as the population activity decreases, with the upper bound approximating the median of the tuning widths in the population (dotted lines), although with some undershoot for broader tuning (e.g. black curve).

Mathematical analysis of these more biologically realistic population codes is not trivial, but we have set out in the Appendix some arguments as to why the limiting behavior of the model at low gains should not be strongly altered by the presence of tuning heterogeneity or noise correlations. One caveat is that these arguments address only the asymptotic predictions at infinitesimally small activity levels (i.e. the “endpoints” the curves are approaching at the far right of each plot), and so provide only a partial explanation of the overall strong similarity between the functions plotted in left- and right-hand panels of Fig 2.

Considering next the mixing parameter, which determines the von Mises proportion in the normal-plus-uniform fit, we observe a monotonic decline with decreasing activity (left to right in Figs 2e & f) for both idealized and biological populations. This parameter depends less on the tuning widths (different colored lines) although there is some influence for middling levels of gain particularly in the biological population. In the conventional interpretation, a decrease in this mixing parameter is interpreted as an increase in the frequency of random or guessing responses. The zero-spike state for the idealized population can be considered a guessing state, although the von Mises proportion of the normal-plus-uniform fit only approximately tracks the probability of zero spikes in the model. However, this possibility of obtaining no information about the stimulus is a unique consequence of the artificial homogeneity imposed on the idealized population. Even the smallest deviation from uniformity in the density of tuning functions over the stimulus space will make zero spikes a more probable outcome for some stimulus values than others, with the consequence that even a zero-spike state would convey some information about the stimulus. In fact, for reasons of computational efficiency, the heterogeneous simulations modeled variation in neural activity with a continuous multivariate Gaussian distribution, rather than a discrete Poisson distribution, demonstrating that these results do not actually depend on the discrete nature of spiking. The normal-plus-uniform fit nonetheless ascribes a large proportion of responses to the uniform (“guessing”) component, particularly at low gains (Fig 2f).

The results of our simulation studies imply that the normal-plus-uniform model parameters can be reinterpreted from a physiological perspective. We observed that changing the summed activation of the underlying population induces a continuous change in both parameters, but importantly, the amplitudes of the change in each parameter are asymmetric. In particular, at the lower end of the gain spectrum, decreases in activity drive the mixing proportion towards zero, while the s.d. parameter saturates toward an upper bound determined by the tuning widths of the component neurons encoding the stimulus. We next examine whether these predictions made by the population coding model can be empirically corroborated.

### Comparing model predictions to orientation recall data

Our simulations indicate that, for errors generated by decoding feature information from a noisy populations of tuned neurons, fits of the normal-plus-uniform model will be bounded, such that the von Mises s.d. parameter cannot exceed the average s.d. of the underlying neural tuning curves even as the precision of the decoded estimates falls to zero. Previous work (Bays, 2014; Taylor & Bays, 2018) has shown that, for multi-item working memory tasks, setting the gain inversely proportional to the number of items in a display provides a good fit of the population coding model to empirical error distributions. On this basis, we should expect the s.d. parameter of a normal-plus-uniform fit to recall data to increase towards an asymptotic value as the number of items increases. Based on the simulations we also predict a monotonic decrease in the von Mises mixing parameter with increasing set size.

Fig. 3 presents normal-plus-uniform parameter estimates obtained from seven previous studies testing recall of orientation stimuli with varying set size. Although there is some variation between experiments, the majority display a common pattern whereby the s.d of the fitted von Mises component increases with set size and begins to saturate (asymptote) when the number of items is large (Fig 3, top). No set size in any study produced a mean von Mises s.d. parameter greater than 20° (out of a 180° space of orientations). This effect on s.d. is paired with a continuous decline in the von Mises mixing parameter with set size (Fig 3, bottom).

**Figure 3.**
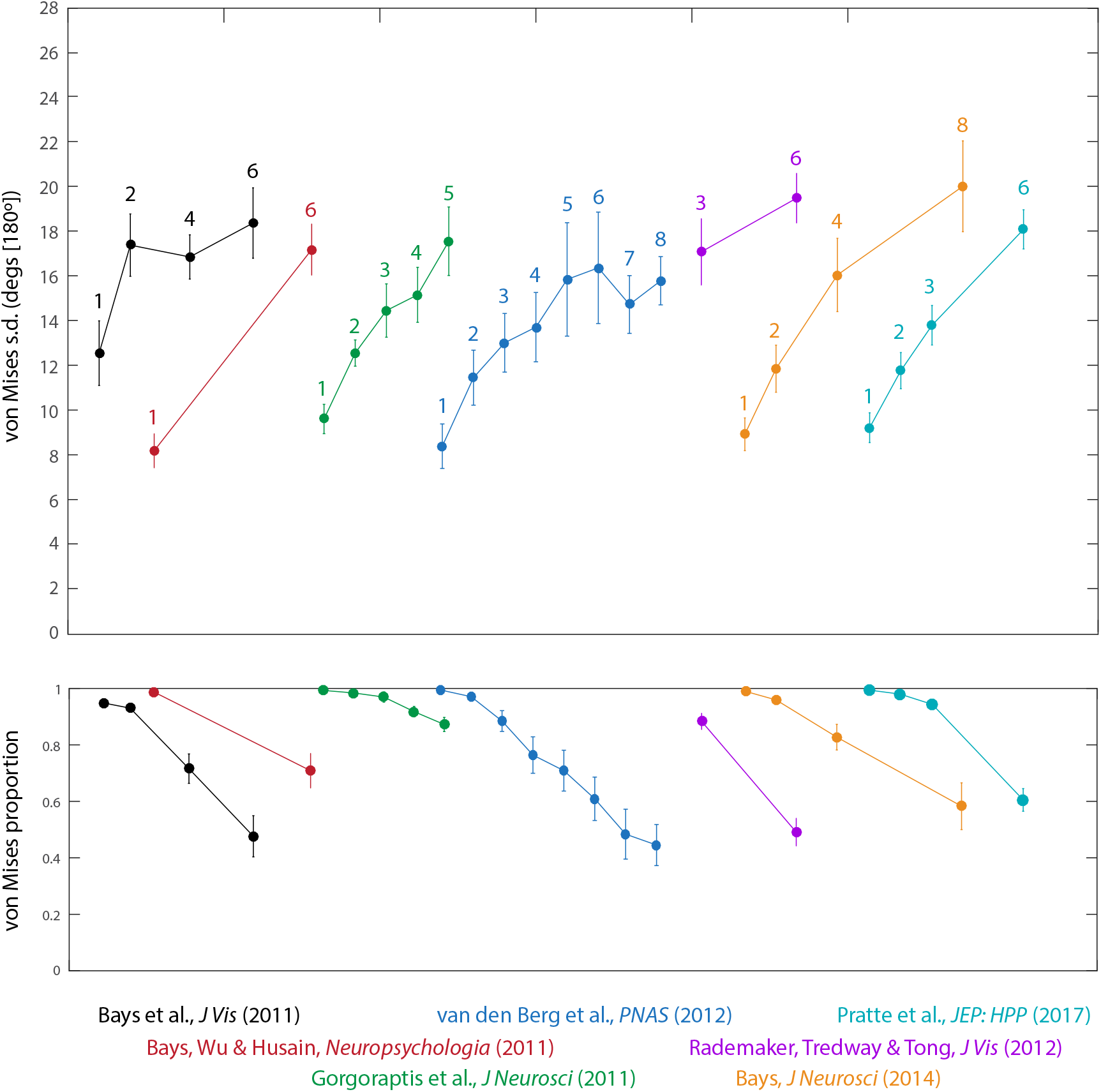
Normal-plus-uniform model parameters estimated from eight orientation recall datasets. The top panel shows the participant-averaged von Mises s.d. parameter obtained across different experiments (colors) and set sizes (numbered). The bottom panel displays the corresponding values of the von Mises mixing parameter. Error bars indicate ±1 s.e. across participants.

To better assess the asymptotic bound on the von Mises s.d. in these studies, we collapsed the data across experiments at the participant level. Fig. 4a plots the results for the von Mises s.d. parameter. The number of participants per set size depends on which conditions were included in each study and is therefore unbalanced: this information is summarized by the relative size of each data point in the plot. With the exception of a single set size (7 items) for which minimal data was available, the mean effect of set size on von Mises s.d. was very accurately fit by a saturating function (red curve; e.g., Albrecht & Hamilton, 1982; Bays, 2018b) with an asymptote at 18.3° [180°] (dashed red line). Results of collapsing the estimated mixture proportion across studies are shown in Fig. 4b.

**Figure 4.**
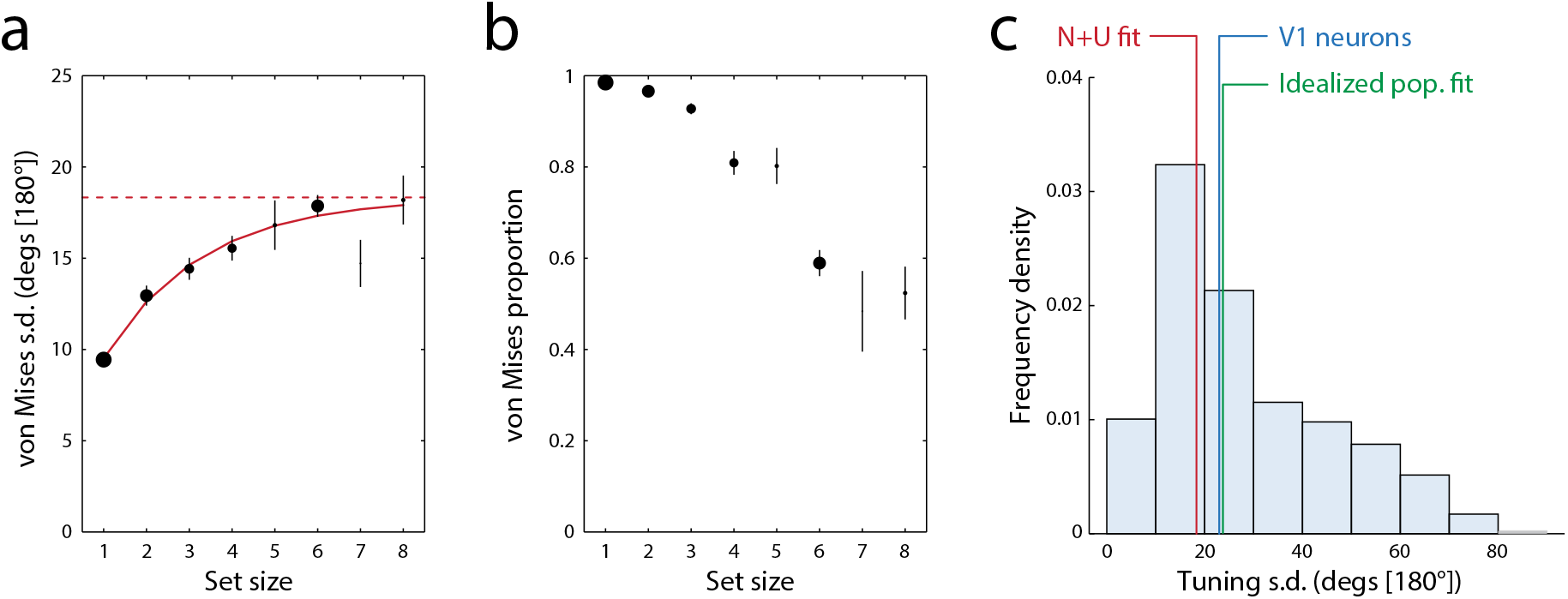
Parameter estimates from combined orientation data. (a) Participant-averaged von Mises s.d. parameter as a function of set size, collapsed across studies. The solid red curve indicates the best fitting saturation function, which has an asymptotic maximum at the level shown by the red dashed line. Size (area) of each data point corresponds to the relative number of participants contributing to it: larger points indicate more data. Error bars indicate ±1 s.e. across pooled participants (b) Participant-averaged von Mises mixing proportions, collapsed across studies. (c) Bars show the distribution of tuning widths (s.d.) recorded from orientation-selective neurons in macaque V1 by Ecker et al. (2010); blue vertical line indicates the median of the recorded widths. Red vertical line indicates the asymptotic limit of the von Mises s.d. parameter shown in (a); green vertical line indicates the median tuning width obtained by fitting the idealized population model to the same behavioral data.

These patterns are consistent with the population coding account (Bays, 2014): if the total activity is normalized, larger set sizes will reduce the amplitude of the neural signal on which individual estimates of feature values are based, which in turn results in larger recall errors. This increase in variability is reflected in complementary shifts in both mixture model parameters: lower activity causes the s.d. of the fitted von Mises component to increase, but also reduces the proportion of responses captured by that component. The approach of the s.d. parameter toward an upper bound means that further decreases in gain – for example, due to testing with an even larger set size – will be reflected primarily in changes to the von Mises proportion. In the next section, we evaluate the prediction that the asymptotic bound corresponds with physiological measures of orientation tuning.

### Comparison with neurophysiology

The above results are strongly consistent with our predictions based on simulations of the population coding model, and in particular suggest that, for orientation recall, there is an asymptotic upper limit on the normal-plus-uniform s.d. parameter at roughly 20° [180°]. If our account is correct, this value is indicative of the average tuning width of the neural populations underlying orientation recall in human participants. Fitting the idealized population coding model directly to the orientation recall data corroborated this estimate, resulting in a median tuning s.d. of 23.2° [180°].

Fig. 4c plots the distribution of tuning widths obtained from the 408 neurons recorded in macaque primary visual cortex by Ecker et al. (2010). While there is considerable variation in tuning across cells, the median tuning s.d. of 23.0° [180°] (blue vertical line) corresponds well with both the asymptote obtained from the normal-plus-uniform fit to behavioral data (red vertical line) and the estimate obtained from fitting the idealized model (green vertical line). This result is further corroborated by an older electrophysiological study by De Valois, Yund, and Hepler (1982) that reported a median orientation full-width at half-maximum of 40°, based on 387 neurons also recorded in macaque V1, corresponding to an s.d. of 18.7° [180°].

### Re-evaluating previous results of the normal-plus-uniform method

The evidence described above for a ceiling on attainable values of the normal-plus-uniform s.d. parameter suggests the need for a re-evaluation of previous results obtained with this method. Fig. 5 displays fitted parameters from four previous studies that applied the normal-plus-uniform method to data obtained under varying conditions with the intention of addressing several distinct research questions. These studies all tested recall of colors chosen from a color wheel (defined as a circle in CIE LAB color space). The dashed black line in Fig. 5 (top) corresponds to the average of the maximum s.d. values obtained in each study (although there is neurophysiological evidence for color tuning in visual cortex e.g., Sanada, Namima, & Komatsu, 2016; Conway & Tsao, 2009, the limited availability of single-neuron data for color tuning widths, and difficulty mapping between color spaces, means we do not have a prediction for this bound). The results suggest an upper limit at approximately 22° [360°]. Note that caution is needed in interpreting the apparent similarity between this value and the one obtained for orientation recall data above: the range of orientations covers 180° whereas the color wheel covers 360°, so as a fraction of the parameter space, orientation recall is approximately twice as variable as color recall.

**Figure 5.**
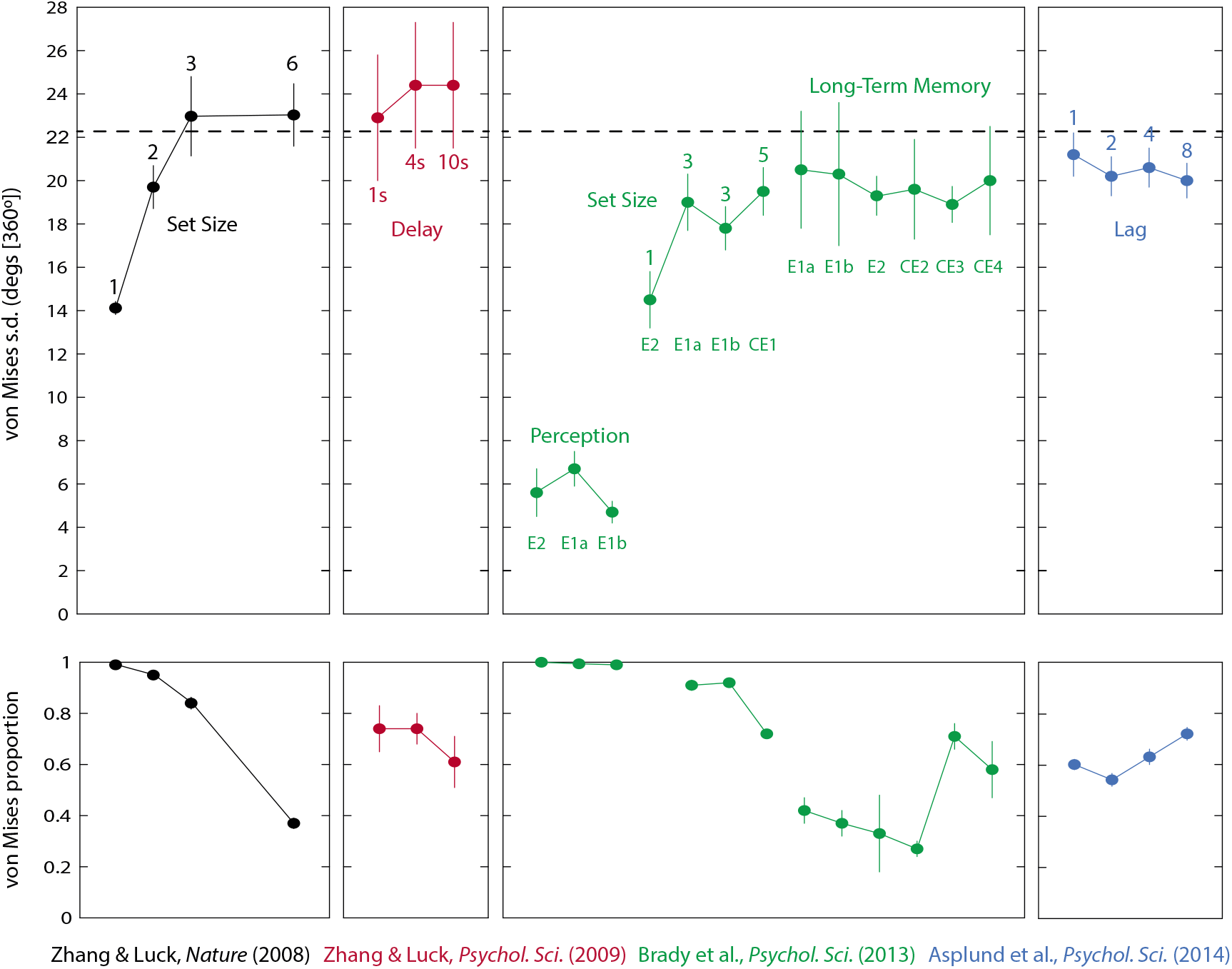
Mixture model parameters obtained in four previous studies using colored stimuli. The top panel shows the participant-averaged standard deviation of the von Mises component for different experimental conditions within each study. The dashed line corresponds to the average of the maximum von Mises s.d. in each study. The bottom panel displays the corresponding participant-averaged von Mises mixture proportions. Note that there is one data point (mixing parameter for set size 1) that was not available from the original paper. Error bars indicate ±1 s.e. across participants with the exception of Zhang & Luck (2009) where they indicate within-participant 95% confidence intervals (as in the original paper).

#### Color Reproduction

Results from W. Zhang and Luck (2008) are shown in black in Fig 5 (far left). This was the first working memory study to fit data with the normal-plus-uniform method, and the sharp plateau in the von Mises s.d. parameter at three items was presented as evidence for an upper limit on the number of items that can be stored in visual working memory. Specifically, assuming that the value of this parameter was a measure of precision for items stored in memory, the authors argued that for small set sizes, multiple independent copies of a single item could be stored in the brain and averaged at recall to enhance memory precision. This “slots-plus-averaging” model predicts that the s.d. parameter will increase with set size (as the number of copies per item decreases) but will abruptly reach a ceiling at the maximum number of items that can be encoded – i.e., the putative capacity limit – beyond which point each item is represented once or not at all.

The sharp plateau at three items observed by W. Zhang and Luck (2008) has not been widely replicated (e.g. see results of orientation studies in Fig 3; also color results from Bays et al., 2009), but the fact that the von Mises s.d. parameter does not increase indefinitely with number of items is a consistent observation across studies. However, our modeling results point to a new interpretation of this finding: rather than a limit on how many items can be stored, the asymptotic upper bound on the s.d. parameter corresponds to, and is a consequence of, the tuning specificity of the neural populations underlying color recall. According to this interpretation, memory representations continue to increase in variability at higher set sizes, but the normal-plus-uniform fit captures this increasing variance primarily with decreases in the von Mises mixing parameter (Fig. 5, bottom) rather than further increases in the von Mises s.d. parameter. Consistent with this interpretation, previous studies that directly fit the population coding model to trial-by-trial responses have found it to provide a consistently better fit than the “slots-plus-averaging” model (Bays, 2014; Taylor & Bays, 2018).

#### Color reproduction with variable retention intervals

A follow-up study (W. Zhang & Luck, 2009) held set size constant at three items, but varied the retention interval between 1 and 10 s. Results obtained from the normal-plus-uniform method are shown as red points in Fig. 5 (center left). The observation of a decrease in the von Mises mixing proportion at the longest delays, coupled with relatively small increases in the von Mises s.d. parameter with delay, were interpreted as evidence for “sudden death”: rather than becoming gradually less precise over time, it was proposed that items spontaneously disappear from memory. The present results again suggest an alternative interpretation: with a set size of three the s.d. parameter of the normal-plus-uniform fit was already close to its asymptotic limit with a 1 s delay (compare with the highest set sizes in data from W. Zhang & Luck, 2008; black points). Further increases in variability at longer delays were therefore captured primarily by decreases in the mixing parameter rather than increases in the s.d. parameter, as predicted by the population coding model.

In the study of W. Zhang and Luck (2009), small increases in the s.d. parameter with delay were observed but did not attain statistical significance. Proposing that this could be a consequence of insufficient statistical power, a recent empirical study (Rademaker, Park, Sack, & Tong, 2018) revisited their design but asked participants to hold just one item in memory over variable delays. Their observation of a reliable increase in the s.d. parameter as delay increased is predicted by our model: the drop in set size to one item shifts the s.d. parameter away from its upper bound, into a regime where changes in precision are reflected in both parameters of the mixture model. More generally, it is important to emphasize that the population coding model does not predict a “plateau” of the s.d. parameter as a result of any of the manipulations considered in this paper, where we take this term to mean a discontinuity in the first derivative of a function, such that its rate of rise abruptly falls to zero. Instead the s.d. parameter is predicted to “saturate” as population gain decreases, i.e. progressively approach an asymptotic upper bound that would be attained only if activity fell precisely to zero (at which point the von Mises contribution to errors would be zero). As a consequence, it should always be possible to detect changes in the s.d. parameter in an experiment with sufficient statistical power. Note that the same consideration applies to the von Mises mixing proportion, which saturates to unity at high levels of population activity.

#### Long-term memory

The green data points in Fig 5 (center right) plot results from a study comparing working memory fidelity to perceptual and long-term memory (Brady et al., 2013). The authors interpreted the apparent upper limit on von Mises s.d. as evidence that items stored in long-term memory share the same fidelity as items held in visual working memory: specifically, this was based on the similarity in this parameter between their long-term memory task and the largest set size (five) in their working memory task. It should be noted that the von Mises mixing parameter drops markedly between the two tasks (Fig 5, bottom), which would conventionally be interpreted as indicating many more retrieval failures in the long-term memory task. In light of our present results, however, this pattern in the mixture model parameters has a simpler interpretation: variability in both cases is sufficient to bring the von Mises s.d. close to its upper bound. This suggests that retrieving a single item from long-term memory into working memory provides only a weak signal: weaker even than the signal from a working memory display with five items. Because five items is already sufficient to bring the von Mises s.d. close to the upper bound determined by neural tuning, this further decrease in signal strength in the long-term memory task is primarily reflected in the von Mises mixture proportion rather than the s.d. parameter.

A recent conceptual replication of the Brady et al. (2013) study, using a larger number of participants and a Bayesian analysis, found evidence for a larger von Mises s.d. in recall from long-term memory than on a three-item working memory task (Biderman, Luria, Teodorescu, Hajaj, & Goshen-Gottstein, 2018). Like the Rademaker et al. study above, this finding is consistent with the predictions of our model: a larger sample size, coupled with a lower set size, means more statistical power to detect small deviations of the s.d. parameter from its asymptotic maximum.

#### Attentional blink

Finally we examined the results of a study by Asplund et al. (2014) using a variation on the classic “attentional blink” task (Shapiro, Raymond, & Arnell, 1997). In this study, participants were presented with a series of colored circles in rapid sequence. Embedded within each stream were two square stimuli, and participants had to report the color of each. The data shown as blue points in Fig 5 (far right) is for a normal-plus-uniform fit to errors in recall of the second target, as a function of the lag between targets (1, 2, 4 or 8 serial positions). Based on these parameters, the authors concluded that conscious awareness of a target item arises in an all-or-none fashion, because the interfering effect of the first target on the second was observed as changes in the von Mises mixing proportion, but not the von Mises s.d. parameter. Again, however, we can see that the von Mises s.d., even at the longest lags where interference should be negligible, is close to the upper bound (dashed line) inferred from comparison across studies^1^. As a result, the observation that the mixing parameter is most strongly affected by lag means only that signal strength is attenuated for the second target when its presentation overlaps with processing of the first, and is not evidence for all-or-none storage.

## Discussion

Thanks to its intuitiveness and ease of implementation, the normal-plus-uniform method has become a common tool for analyzing continuous report data. Indeed, the ability to extract from response errors two parameters related to performance – instead of a single measure of dispersion – can seem like a way of getting “something-for-nothing” from an experimental dataset. The interpretation of these parameters as products of separable psychological processes determining the quality and quantity of representation has led researchers to make claims of “discreteness” not only for working memory (W. Zhang & Luck, 2008) but also iconic (sensory) memory (Pratte, 2018) and even conscious visual perception (Thibault, van den Berg, Cavanagh, & Sergent, 2016; Asplund et al., 2014).

Yet, the standard interpretation placed on normal-plus-uniform parameters can in some cases produce surprising, even incompatible, conclusions. Consider the “bilateral field advantage”, the observation that memory is improved for visual items that are distributed across both the left and right visual fields, compared to within a single hemifield (Delvenne, 2005). Using the normal-plus-uniform model, Umemoto, Drew, Ester, and Awh (2010) came to the conclusion that this bilateral advantage is due to an increase in the number, but not the precision, of stored items. However, with minor modifications to the timing and number of items, Y. Zhang et al. (2017) subsequently found that the advantage manifested as a change in precision, with no effect on storage probability.

Another example has arisen in the application of the normal-plus-uniform method to investigation of mental disorders. Based on a color report task, Gold et al. (2010) concluded that, though patients with schizophrenia stored items just as precisely as healthy controls, they tended to store fewer items in memory at one time. In comparison, Xie et al. (2018), using the same task, reported instead that individuals with self-reported schizotypy remembered items less precisely and did not differ in how many items were remembered. The fact that small changes in experimental design, or testing a different population drawn from the same diagnostic spectrum, can lead to such conflicting conclusions is troubling for the psychological model. But when the representation of visual items in neural activity is taken into consideration, these findings become much less surprising, because we have shown that both components of the normal-plus-uniform model are influenced by changes in the same underlying parameter, i.e. the amplitude of the neural signal encoding the information.

Recently, Bays and Taylor (2018) set out to resolve similarly conflicting results from studies of retrospective cuing. It is well established that recall performance for an item can be enhanced by directing attention to its representation in memory, even after the stimulus itself has disappeared (Griffin & Nobre, 2003; Souza & Oberauer, 2016). However, the normal-plus-uniform method has largely proved uninformative about the nature of this effect: retro-cueing is associated with both a decrease in the von Mises s.d. parameter and an increase in the von Mises mixture parameter (as well as a decrease in swap errors). In comparison, when Bays and Taylor (2018) fit the population coding model to data from previous retro-cue experiments, they found the effect of the retro-cue was expressed as a simple increase in the neural gain, with no change in tuning width. This result parallels that found for prospective cues that orient attention to an item’s location before it appears (Bays, 2014). A decrease in swap errors can also be explained by increased gain (Schneegans & Bays, 2017), so the population coding model effectively reduces three behavioral effects down to a single underlying cause.

In this study we examined the consequences of fitting the normal-plus-uniform mixture model to error distributions resulting from decoding features represented in noisy neural activity. The results of our simulation studies (Fig. 2) demonstrated the following qualitative features. When the population activity level (gain) was high, any decrease in gain was primarily observed as an increase in the von Mises s.d. parameter, whereas the von Mises mixing proportion was relatively unchanged at about one. Conversely, if the gain was already low, then further decreases in gain were reflected mainly in decreases of the mixing proportion, while the von Mises s.d. parameter asymptotically approached a ceiling value equal to the average tuning width of the underlying neural population. This asymptotic behavior could be predicted based on a mathematical analysis of an idealized neural population, and critically, this pattern was found to be largely robust to variations in the implementation of the model intended to more closely approximate the heterogeneous properties of visual neurons recorded *in vivo*.

Based on existing behavioral results (Bays, 2014; Taylor & Bays, 2018) and neurophysiological evidence suggesting that the neural activity encoding individual stimuli in memory decreases with set size (Buschman, Siegel, Roy, & Miller, 2011; Sprague, Ester, & Serences, 2014), we predicted that the asymptotic bound on von Mises s.d. would be observed in tasks in which the number of items in memory was manipulated. Compiling data from numerous previous studies of working memory for oriented stimuli, we found evidence for such an upper bound at approximately 20° of orientation (in a 180° range). According to our model, this bound is indicative of the average tuning width in the neural populations underlying orientation memory, and indeed the observed value compares rather accurately with the median orientation tuning width of cells recorded in macaque primary visual cortex.

This finding should be treated with some caution, not least because the behavioral results and the recordings come from different species. While the anatomy of primary visual cortex is reasonably well conserved across primate species (Rosa & Tweedale, 2005), we are not aware of any systematic comparison of visual tuning properties between humans and macaques. Indeed we know of no single cell recording studies of human visual cortex, and current methods are unable to extract tuning widths from non-invasive techniques such as fMRI or EEG (Sprague et al., 2018). A study of single neuron responses in human auditory cortex found that frequency tuning was more narrow than is typical for mammals, including macaques (Bitterman, Mukamel, Malach, Fried, & Nelken, 2008).

We chose orientation tuning in area V1 as the basis of our neurophysiological comparison primarily because of the availability of suitable electrophysiological data. Our model is agnostic as to brain region, and the kind of population coding on which it is based is observed widely in the brain. Whether the intrinsic properties and connectivity of V1 make it a viable candidate region for maintaining memory representations is debated (Serences, 2016; Xu, 2017; Harrison & Bays, 2018; Bloem, Watanabe, Kibbe, & Ling, 2018; Rademaker, Chunharas, & Serences, 2019). Imaging studies have demonstrated that orientations, locations and other visual features maintained in working memory can be decoded from signals originating in multiple brain areas within occipital, parietal and prefrontal cortex (Sprague et al., 2014; Christophel, Iamsh-chinina, Yan, Allefeld, & Haynes, 2018).

To what extent are the V1 tuning parameters that were our focus here representative of visual areas more widely? Only a very few studies have directly compared orientation tuning of cells in primate V1 to cells in other visual areas, and comparison across studies is made complicated by differences in the methods by which tuning was assessed. Nonetheless, the available evidence indicates tuning widths do not vary greatly between areas, e.g. Gegenfurtner, Kiper, and Levitt (1997) reported a median half-width at half-maximum (HWHM) of 27.2° in V3, only 14% higher than the 23.9° in the Ecker et al. V1 data; that paper further reported two previous studies that obtained very similar median HWHMs in V2: 26.7° (Gegenfurtner, Kiper, & Fenstemaker, 1996) and 29.7° (Levitt, Kiper, & Movshon, 1994). McAdams and Maunsell (1999) made a direct comparison of averaged tuning functions recorded from 197 neurons in V4 and 125 neurons in V1, obtaining widths that differed by < 1%. Albright (1984) found that MT neurons sensitive to the orientation of stationary slits had tuning approximately 23% wider than V1. Assessing orientation tuning widths at higher levels of the visual hierarchy becomes increasingly challenging as few studies have systematically measured tuning for more than a handful of cells, and also because neurons typically have responses that depend in a non-linear fashion on the conjunction of multiple visual features. However, we have seen no evidence to suggest tuning of those neurons with orientation selectivity in, e.g. area IT (Tanaka, Saito, Fukada, & Moriya, 1991), is significantly broader than V1.

A final consideration is that there is substantial inter-neuron variability in tuning width in all areas that have been systematically assessed. This makes it difficult to obtain an unambiguous estimate of the average width, a problem compounded by the possibility that very broadly-tuned neurons may not have met a study’s inclusion criteria for orientation selectivity.

In attempting to relate a behavioral characteristic, obtained from patterns of errors in orientation reproduction tasks, to a property of the neural system, specifically the tuning of orientation-selective visual neurons, we are conscious that we are making what some might consider a bold claim. The population coding account of working memory was developed with the intention of capturing general principles of neural representation, and demonstrating that they could explain aspects of the distribution of behavioral reproduction errors that were otherwise difficult to account for. It is certainly a strong version of this hypothesis that predicts a direct correspondence between these errors and tuning parameters of samples of visual neurons recorded *in vivo*. Nonetheless, the prediction for neurophysiology is clear, and the present results indicate that the strong population coding hypothesis has indeed survived this first test.

On a more pragmatic level, and irrespective of a direct correspondence to neurophysiology, our results indicate a need to reconsider the usual interpretation of normal-plus-uniform model parameters, and to re-evaluate previous conclusions based on observed changes in those parameters. In particular, the pattern of results in which an experimental manipulation decreases the von Mises mixture proportion, but leaves the von Mises s.d. unchanged, has been used to argue for a fixed item limit on working memory capacity (W. Zhang & Luck, 2008), that memories experience “sudden death” rather than gradual decay (W. Zhang & Luck, 2009), that short-term and long-term memory have a common limit on fidelity (Brady et al., 2013), and that conscious awareness is an all-or-nothing process (Asplund et al., 2014), among other examples.

In contrast, our results indicate that this pattern of results can arise simply as a consequence of model mismatch: fitting the normal-plus-uniform model to the distributions of error predicted by population decoding produces the same pattern of changes under conditions of low neural gain, even when the neural model contains no element of discreteness and no guessing process (as in the case of simulated heterogeneous populations, which we believe most accurately reflect cortical neurophysiology).

The interpretation of the uniform component in working memory tasks as solely due to random guessing has been criticized previously, on the grounds that many of the responses captured by this component are not random, but are in fact systematically distributed around the feature values of other items in memory (Oberauer & Lin, 2017; Schneegans & Bays, 2017). It has also been shown (Bays, 2018a) that estimates of the putative upper limit on working memory capacity derived from the mixing parameter do not coincide with estimates of the upper limit derived from the s.d. parameter, contrary to predictions of memory models with fixed limits (W. Zhang & Luck, 2008). However, the present results go further, by demonstrating that changes in the mixing proportion of the normal-plus-uniform model could arise from a simple change in the amplitude of neural signal underlying a memory, and conversely that no change in the von Mises s.d. parameter may be observed even when an experimental manipulation decreases the precision of representation.

Although we question the ability of the normal-plus-uniform fit to index guessing responses, and consider the hypothesis of a fixed upper limit on the number of objects in working memory to be largely refuted, it is not our contention that guesses do not occur. At a practical level, many of the studies examined here did not track eye movements, meaning that on some trials participants may have been looking elsewhere, or blinking, during presentation of the memory array, inevitably resulting in responses that are randomly distributed relative to the memoranda.

On the remaining trials, the relevant visual information will have been detected by photoreceptors in the participant’s retinas and transduced into neural activity, altering the state of the neural system. At this stage defining a guess becomes more challenging. For a brain state to convey zero information about a stimulus, the probability of that brain state arising must be identical for every possible stimulus value: even in a hypothetical situation where an attempt to recall a stimulus elicited no spikes from an encoding population, the very fact that no spikes were emitted would likely provide evidence favoring one value for the stimulus over another. However, zero information may be too strict a criterion on the concept of guessing. A key aspect of resource-based models of working memory, distinguishing them from the older “slot” models, is that recall responses exist on a spectrum from high-confidence accurate reproductions of the target to low-confidence reports with large errors that could reasonably be described as “guesses” (although without requiring a separate guessing mechanism). The population coding model provides a putative neural basis for why responses exist along a continuum. As a result, it has been successful in modeling meta-cognitive data, reliably reproducing patterns of confidence judgments and the relationship between rated confidence and recall precision (Bays, 2016b).

Our claim is that the two parameters of the normal-plus-uniform mixture model do not correspond to two independent neural or psychological processes of response generation. Instead, both are driven by the strength of the memory signal, corresponding to the amplitude of the underlying neural activity. An unpublished manuscript by Schurgin, Wixted, Brady, et al. (2018) provides a complementary interpretation of normal-plus-uniform parameters. These authors similarly argue that the s.d. and mixture proportion parameters do not reflect separable processes but are instead both driven by a single memory strength signal. They propose that the long-tails of recall error distributions are the result of a non-uniform transformation between the physical metric space on which experimental stimuli are defined (e.g. a color wheel) and the internal psychophysical metric space in which they are perceived or stored. On examination, these authors’ Target Confusability Model was found to implement a form of winner-take-all population decoding, with a remarkably close correspondence to the neural models addressed in the present study. For detailed discussion see Bays (2019).

It is worth noting that some of the arguments above find parallels in the long-term memory literature, where they have been deployed against high-threshold models of recognition memory and in favor of signal detection accounts based on a continuous-valued familiarity signal (e.g. Wixted, 2007). The observation of similar patterns of errors in reporting features retrieved from long-term memory as working memory (Schurgin et al., 2018; Brady et al., 2013) is predictable based on the population coding account, which does not require that features are maintained in the form of persistent spiking during the memory delay, but only assumes that they are represented in spikes at retrieval. In particular, it is reasonable to assume that a feature retrieved from a passive store such as long-term memory will need to be actively represented in spiking activity in order to be reported or reproduced (in cognitive terms, it must be brought into working memory). We therefore expect the same constraints on error distributions, imposed by the tuning parameters of the active neural population, to apply as in a working memory task.

It is important to acknowledge that the normal-plus-uniform model has value as an analytical tool for psychological and neuroscientific research. For numerous reasons, including but not limited to the aforementioned blinks, gaze deviations and swap errors, data collection may yield a set of observations that contain contaminants arising from processes other than those of interest. In many cases it may be desirable to remove these contaminant trials even if that means removing some valid data points as well, and the normal-plus-uniform model (or the extended three-component model; Bays et al., 2009) can provide a valuable statistical tool for data pre-processing, as long as the limitations of the approach are recognized. To reiterate, any attempt to interpret the fitted s.d. of only the normal component of error as a measure of representational precision must bear in mind it is a bounded estimate, with the consequence that an absence of change in this parameter is not evidence for fixed precision, unless one can also show the parameter value is well below its upper bound. Additionally, responses captured by the uniform component of the model can arise from a number of different processes – including imprecise recall of the target feature, swap errors, or basing responses on hierarchical representations or ensemble statistics (Brady, Konkle, & Alvarez, 2011, 2009) – and should not be interpreted as a simple measure of guessing. The fact that these influences are primarily absorbed by the uniform component of the fit may in part be why an effect of the underlying tuning can be recovered.

It will be important to corroborate the evidence presented here for a relationship between behavioral performance on continuous recall tasks and tuning characteristics of the underlying neural populations, in particular using a range of different feature dimensions, and comparing behavior to neurophysiology within individuals. However there are practical difficulties in doing so, both because acquiring tuning functions from a sufficient sample of neurons is an arduous and invasive neurophysiological task, rarely attempted in humans, and because the continuous report task is poorly suited to the requirements of training and testing non-human species. We note some recent advances that suggest these challenges may soon be overcome, first through the use of two-photon neuroimaging to simultaneously record individual tuning of large numbers of visual neurons (Ikezoe, Amano, Nishimoto, & Fujita, 2018), and second in reports of the first multiple-item continuous recall data from non-human primates (Panichello, DePasquale, Pillow, & Buschman, 2019).

## Appendix

### Idealized neural populations

We begin by considering a population of *M* idealized neurons encoding a stimulus characterized by an angle, *θ*, where the tuning functions of each neuron *f*(·) are unimodal and identical except for translation around the circle, such that preferred values (the modes of the tuning functions) are distributed uniformly throughout the angular space [-*π*, *π*). The mean response of the *i^th^* neuron with preferred value *φ_i_* to stimulus *θ* is:

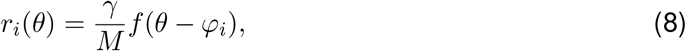

where *γ* is the population gain, i.e. total expected activity summed over the whole population of neurons.

We model spiking activity of each neuron as a stationary (i.e. time-invariant) Poisson process, uncorrelated with the activity of other neurons, such that the probability of the *i*th neuron generating *n* spikes during a decoding interval of length *T* is:

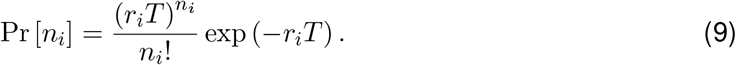

Retrieval of the encoded stimulus value is based on maximum likelihood (ML) decoding of the population’s joint spiking activity **n**,

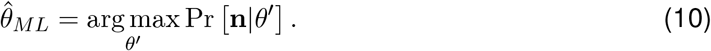

Because neurons are statistically independent, the total number of spikes available for decoding is also Poisson distributed,

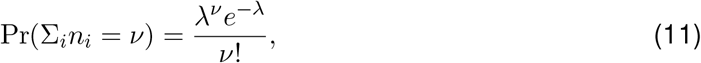

where λ = *T*Σ*_i_r_i_*, which, assuming the population provides a dense uniform coverage over the stimulus space, is equal to *γT*.

We can write the distribution function for the decoded estimate 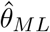 as a power series,

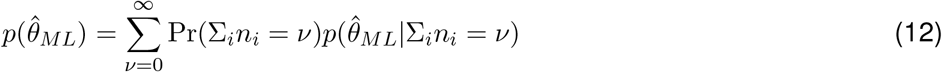

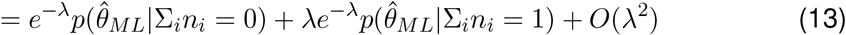

where *O*(λ^2^) indicates terms of order λ to the power of two or greater (for λ → 0).

It follows that, as the population gain *γ* tends to zero, and hence λ also tends to zero, the distribution of the decoded estimate will tend towards a mixture of two distributions, corresponding to population spike counts of zero and one. The former distribution, 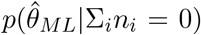, will be uniform, because if there are no spikes, Pr[**n**] is independent of *θ*. The distribution of responses for a spike count of one can be obtained by expanding and taking the logarithm of Eq. 10,

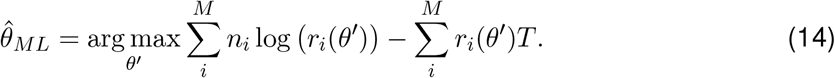

Assuming, again, a dense uniform coverage, the second term is constant and can be ignored. As we are considering the case that only a single neuron (the *j^th^*) generates a spike, i.e. *n_j_* = 1, *n_i≠j_* = 0, the decoded estimate is,

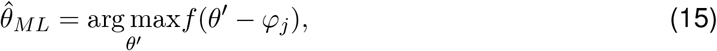

which is, by definition, equal to the preferred value *φ_j_* of the neuron that generated the spike. The probability that this is the *i^th^* neuron is proportional to *r_i_*(*θ*). Based on Eq. 8, and assuming dense uniform coverage, we find that,

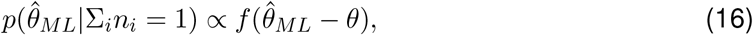

i.e. the decoded estimate is distributed around the true stimulus value with probability density equal to the (normalized, reflected) tuning function. Combining this with the result for zero spikes, we conclude that as gain *γ* → 0 the distribution of errors will tend towards a mixture of a uniform distribution and a distribution matching the tuning function. As a case in point, if the tuning function is von Mises with concentration parameter *κ*, then the error will be distributed as a mixture of a uniform distribution and a von Mises distribution with the same concentration *κ*. This corresponds to the result we obtained from simulation (left-hand panels of Fig 1 in main paper).

### Biological neural populations

The responses of real neurons that exhibit population coding, such as orientation-selective neurons in V1, differ in a number of ways from the idealized population considered above. Nonetheless, the results of numerical simulations of populations with more plausible tuning properties (e.g. based on the tuning heterogeneity of neurons recorded in V1; right-hand panels of Fig 1 in main paper) show a remarkable similarity in their pattern of decoding errors to the idealized case. Here we address each of the most salient differences between real and idealized populations in turn and consider to what extent the results obtained above can be expected still to hold. While the following falls short of a full mathematical treatment, we believe it is useful in providing intuitions for the numerical simulation results reported in the main paper.

#### (a) Baseline activity

Most neurons in real populations do not fall entirely silent even when stimulus features differ strongly from the neurons’ preferred values: their tuning functions can be approximated by the sum of a von Mises function and a constant corresponding to a baseline level of activity. The consequences for error distribution at low gains are already accounted for in the equations set out above: one-spike errors will be distributed in correspondence with the tuning function (Eq 16), i.e. as a mixture of von Mises and uniform distributions. Zero-spike errors will still be uniform. So the only observable effect will be that, for a given population gain, the von Mises mixing proportion will be diminished compared to a population without baseline activity (and, as a corollary, recall estimates will be less precise). The width of the von Mises component of the error distribution will have an upper bound at the width of the von Mises component of the tuning function.

#### (b) Non-uniform distribution of preferred values

Real neural populations can show both systematic and unsystematic deviations from uniform spacing of preferred values. As a result the second term in Eq 14, the expected summed activity of the population, will not be constant across changes in the stimulus: certain stimulus values will elicit more activity overall than others. The most significant impact is in the zero-spike case: the absence of firing now provides information about the stimulus, so that rather than being uniformly distributed, estimates will be biased towards particular regions of the parameter space. However, because our interest here is in the distribution of estimates *relative to the true stimulus value*, and – in the experimental tasks we are examining – stimulus values were chosen at random from a uniform distribution, biases induced by non-uniformity of preferred values will have no observable effect on the zero-spike component of errors, which will be uniform just as for the idealized population. Nonuniformity in preferred values will also bias responses based on one or more spikes, although more weakly and with an effect that diminishes the more spikes are available for decoding. Relative to the true stimulus value, these biases are expected to manifest as a broadening of the error distribution, so that as gain *γ* → 0, the width of the non-uniform component of errors will tend to exceed to some degree the width of the tuning functions.

#### (c) Variation in tuning amplitude and width

Neural responses recorded *in vivo* show considerable heterogeneity in specificity and amplitude. Like non-uniformity in preferred values, this will cause variation in the total expected activity (second term in Eq 14) across stimulus values, with the same consequences for decoding as described in (b). Additionally, because errors in the one-spike state are distributed like the tuning function of the neuron that fired the spike, variations in tuning width will make this distribution a scale mixture of von Mises distributions of different concentrations. Because neurons with higher response amplitude are more likely to have generated the spike, the mixture will be weighted by amplitude such that it reflects most strongly the tuning widths of the most responsive neurons. So at low gains, errors will be distributed as a mixture of a uniform distribution and a scale mixture of von Mises distributions. It is difficult to predict the consequences of fitting such a distribution with a mixture of a uniform and a single von Mises distribution, but assuming the variation in tuning is not too great it is likely that the fitted von Mises component width will approximately correspond to the average width of the tuning functions, and indeed this is what we observed in our numerical simulations.

#### (d) Interneuronal noise correlations

Correlations in population activity (specifically, “noise” correlations, i.e. those that are independent of the stimulus) are the most challenging factor to incorporate into the model, primarily because they are the least well characterized by electrophysiological recordings. The most conspicuous are “limited-range” pairwise correlations, the tendency for neurons with similar preferred stimulus values to spike in concert, and these are the type of correlation we examined in our simulation study. The consequences of these correlations for decoding depend on their strength (which has been inconsistently estimated across recording studies, see Cohen & Kohn, 2011; we chose a value at the upper end of empirical estimates for our simulations), and also the extent to which the decoder is informed by the correlation structure in the population: in a heterogeneous population a decoder that “knows” about the correlations can to some extent nullify their effects (Ecker et al., 2011). Recent theoretical work suggests that the most significant correlations with respect to limiting the information content of a population are “differential correlations”, i.e. correlations that match those induced by small random changes in the stimulus value (Moreno-Bote et al., 2014). However, the strength of these correlations in real neural populations is difficult to establish (Kohn, Coen-Cagli, Kanitscheider, & Pouget, 2016).

Despite these uncertainties, there is reason to believe the presence of correlations will not strongly influence the patterns of decoding error at low gains. First, while correlations violate the assumption of independence between neurons used to derive Eq 11 – meaning that summed activity may not be Poisson distributed – so long as there are no perfectly correlated neurons (i.e. all correlation coefficients are less than one) the probability of obtaining *n* spikes will still decrease in proportion to λ^*n*^ as λ → 0 (as in Eq 13), so we can still justifiably base our predictions for low gain decoding on the limiting case of one and zero spikes. Because the marginal distributions of each neuron’s spike count are the same as for an uncorrelated population, we do not expect the presence of correlations to have systematic effects on the decoding of a single spike, or no spikes. So while noise correlations have an uncertain and potentially significant effect on the information content of population codes in general, their influence in the limit of low population activity is likely to be minimal.

#### Summary

The simulation results reported in the main paper indicate that the key predictions for error distribution under low gain obtained by mathematical analysis of an idealized neural population hold to a good approximation for populations with tuning properties more representative of the biological system. In the above we have presented some arguments as to why this should hold true, based again on consideration of the limiting case in which decoding relies on one or zero spikes. We emphasize that we are not claiming that decoding of single spikes makes a meaningful contribution to human recall errors; only that investigations of limiting cases can provide important insights into a system’s behavior away from those limits. In particular, we suggest that biological populations – as a consequence of baseline activity, tuning heterogeneity, interneuronal correlations, and possibly other factors not yet considered – may demonstrate at higher activity levels some of the same decoding properties as an idealized population at the very lowest levels.

## Footnotes

1 The supplementary materials of Asplund et al. (2014) report an additional control experiment using a reduced rate of stimulus presentation. The mean von Mises s.d. parameter of ~16° is lower than in the main experiment, making this potentially a better test for a disruptive effect of the first target on this parameter. Unfortunately the results are not decisive: although no significant effect of lag on s.d. was observed, the evidence for an effect on the uniform component is also weak, perhaps because the slower presentation attenuated the attentional blink effect. The experiment was also comparatively low powered (11 participants, compared to 28 in the main experiment) which could explain the null result.

